# Relative brain size and cognitive equivalence in fishes

**DOI:** 10.1101/2021.02.09.430417

**Authors:** Zegni Triki, Mélisande Aellen, Carel van Schaik, Redouan Bshary

## Abstract

There are two well-established facts about vertebrate brains: brains are physiologically costly organs, and both absolute and relative brain size varies greatly between and within the major vertebrate clades. While the costs are relatively clear, scientists struggle to establish how larger brains translate into higher cognitive performance. Part of the challenge is that intuitively larger brains are needed to control larger bodies without any changes in cognitive performance. Therefore, body size needs to be controlled to establish the slope of cognitive equivalence between animals of different sizes. Potentially, intraspecific slopes provide the best available estimate of how an increase in body size translates into an increase in brain size without changes in cognitive performance. Here, we provide the first evaluation of this hypothesis for fishes. First, a dataset of 51 species that included only samples of ≥ ten wild-caught individuals yielded a mean brain-body slope of 0.46 (albeit with a large range of 0.26 to 0.79). This mean slope is similar to the encephalisation quotients for ectotherm higher taxa, i.e. teleost fishes, amphibians and reptiles (∼ 0.5). However, the slope is much higher than what has been found in endotherm vertebrate species (∼ 0.3). Second, we provide slope estimates for brain-body sizes and for cognition-body sizes in wild-caught cleaner fish *Labroides dimidiatus* as a case study. Brain-body slopes from two datasets gave the values of 0.58 (MRI scans data) and 0.47 (dissection data). Furthermore, we have cognitive performance data from 69 individuals tested in four different cognitive tasks that estimated learning, numerical, and inhibitory control abilities. In all four tasks, the cognitive performance did not correlate significantly with body size. These results suggest that the brain-body slopes represent estimates of intraspecific cognitive equivalence for this species. While subject to further studies on various species, our results suggest that endo- and ectotherm brain organisations and resulting cognitive performances are fundamentally different.

## Introduction

Scholars have long sought regularities in the relationship between brain size and body size. These analyses, largely focussing on mammals, showed an unexpected pattern: the slope of (log-transformed) brain size on (log-transformed) body size is often lower within than between species [Pilbeam, and Gould, 1974; Tsuboi et al., 2018; Pagel, and Harvey, 1988]. Two different explanations for this phenomenon, called the taxon-level effect, have been proposed. The first attributes the higher slope at higher taxonomic levels to the combined effect of two parallel macroevolutionary processes: Cope’s rule [Alroy, 1998], the tendency for more recently evolved lineages to have larger bodies than their forebears, and Lartet-Marsh rule [Jerison, 1973; Halley, and Deacon, 2017], the tendency for more recently evolved lineages to have larger brains at a given body than their forebears. The combined effect of the two processes would be to artificially inflate the regression slope through the total sample of all species relative to those within a given species.

A second explanation is based on statistical logic: the estimated slope is reduced when the error in body size measures is greater than that for brain size [see detailed discussion in van Schaik et al., 2021]. This effect is strongly affected by body size range in the sample and thus far greater within species than between them. This phenomenon has been argued to explain why estimates for within-species slopes show extensive variation [Pilbeam, and Gould, 1974], even among different mammalian lineages [Martin, and Harvey, 1985].

While the two explications for the taxon effect should in principle apply to all taxa, Tsuboi et al. [2018] showed that there is a clear pattern in this variation: ectothermic vertebrates (fishes, amphibians and reptiles) have far steeper intra-specific slopes (in their sample ca 0.50) than the endothermic birds and mammals (0.15-0.20). Because the majority of ectotherms show indeterminate growth and thus a far greater range of adult body sizes, the error problem might, in principle, explain this difference. However, Tsuboi et al. [2018] showed that the error problem was unlikely to explain this difference fully. Indeed, a recent analysis focused on sexually dimorphic primates to calculate intraspecific slopes using mean values for males and females, and so minimize the error problem, obtained only slightly larger slopes of 0.25-0.3 [van Schaik et al., 2021]. Nevertheless, it remains unclear what value we should expect for the ectotherms for two reasons. First, the full data set used by Tsuboi et al. [2018] contained both wild and captive fishes, but hatchery-reared fish have smaller brains compared to fish of the same species in the wild [see review by Huntingford, 2004]. Second, they did not analyse how much the range of body sizes in a species’ sample affected the value of the slope. This is important because fishes have a much larger range of body sizes, due to their indeterminate growth, than birds or mammals [Froese, and Froese, 2019]. This feature makes fishes more suitable than birds or mammals to address these statistical issues. Therefore, the first aim of this paper is to refine the findings of Tsuboi et al. [2018] by estimating intra-specific slopes using only wild-caught individuals and assessing the effect of intraspecific variation in adult body size on the estimate. To do so, we identified 51 fish species for which sample sizes of wild specimens were large enough (≥ 10 data points) to calculate brain-body slopes.

Suppose the analyses corroborate the difference between endothermic and ectothermic vertebrates in the slope of the brain-body relationship. In that case, this raises the important question of how the different slopes translate into cognitive performance. We follow Shetlleworth, who defined cognition as the ability to acquire, process, retain information and act on it [Shettleworth, 2010]. This broad definition of cognition may justify analyses on total brain size, while specific areas should be studied when their function is known, like the hippocampus and spatial memory [Krebs et al., 1989]. Regarding total brain size analyses, Jerison [1973] had proposed the encephalization quotient (called EQ), the ratio of a species’ actual brain size to its predicted brain size based on its clade-specific brain-body regression line, to capture its cognitive performance relative to other species within the same clade. The use of EQ as a predictor for relative cognitive performance across species has been criticised largely because of the taxon-level effects reported for endothermic vertebrates, and intraspecific slopes have been proposed as an alternative [van Schaik et al., 2021]. The key idea of this alternative approach is that the intraspecific slope reflects the extra amount of brain tissue required to sustain the additional somatic functions in larger individuals because there are no changes in bauplan and sensory-motor abilities. Therefore, this slope would serve to identify the line of so-called ‘cognitive equivalence’ [van Schaik et al., 2021]. The critical assumption thus is that the cognitive performance of adult individuals within a species is independent of their body size. This assumption can be argued to hold for humans: one good estimate of overall cognitive performance, IQ, shows only a very modest effect of the sex difference in body size [Irwing, and Lynn, 2005]. In addition, cognitive test batteries applied to various species find no effects of body size, including sex differences among dimorphic species [reviewed in van Schaik et al., 2021; see also Bohn et al., 2021]. The markedly higher value of the intraspecific slope in ectotherms than endotherms, therefore, raises the question of whether adult ectotherms of different sizes also show cognitive equivalence or whether larger and hence older individuals outperform smaller ones.

Therefore, the second aim of this study is to provide a first test of the prediction that cognitive performance of adult individuals within a fish species is not correlated with body size (which is also a proxy of age in ectotherms). Although we know little about domain-general intelligence in ectotherms [Aellen et al., 2021, preprint; Poirier et al., 2020], unlike in endotherms [Burkart et al., 2017], we can examine whether a variety of cognitive tests show a body size effect on performance. If they all yield similar results, we can conclude cognitive equivalence in our study species, amenable to further testing in other fish species.

We examined cognitive performance in the cleaner fish *Labroides dimidiatus*, a species for which we also provide the slope of the brain-body relationship. Cleaner fish engage in iterative mutualistic interaction with a variety of coral reef fish species (hereafter “clients”). Cleaners remove client’s ectoparasites and dead tissue in exchange for food [Losey, 1979], a behaviour that is mutually beneficial for both cleaner [Grutter, 1999] and client [Demairé et al., 2020; Ros et al., 2020; Clague et al., 2011; Waldie et al., 2011]. Cleaner fish show a highly sophisticated strategic behaviour that includes reputation management [Bshary, and Grutter, 2002; Binning et al., 2017], social tool use [Bshary et al., 2002], reconciliation [Bshary, and Würth, 2001], and social competence [Triki et al., 2020a]. Cleaner fish may outperform primates in some learning tasks [Salwiczek et al., 2012] that go beyond conditioning [Quiñones AE et al., 2020]. They also show evidence for generalised rule learning [Wismer et al., 2016], numerical discrimination [Triki, and Bshary, 2018], long-term memory [Triki, and Bshary, 2019], and mirror self-recognition [Kohda et al., 2018].

We used published data on cleaner fish cognitive performance in four different cognitive tasks [Aellen et al., 2021 Preprint] to explore the relationship between body size and cognitive performance. These results were then combined with estimates of the brain-body slope in this species using published data from two studies on brain size [Triki et al., 2019a; Triki et al., 2020a]. These data sets also contain information regarding the size of five major brain areas, i.e. telencephalon, diencephalon, midbrain, cerebellum and brainstem. As these areas differ in their relative importance to subparts of cognitive performance (e.g., perception versus decision-making versus execution of actions [Striedter, 2005]), we also explored the regression slopes of the five main brain region sizes on body size in cleaner fish. The brain data and the cognitive performance data were not collected from the same individuals, and thus these datasets are independent. Nevertheless, the results show to what extent intraspecific brain-body slopes in fishes may reflect cognitive equivalence.

## Material and Methods

### Brain and body size data from other fish species

We compiled fish brain and body size data from the database FishBase [Froese, and Pauly, 2019] and the published study by Tsuboi et al. [2018]. We selected only confirmed wild-caught individuals and discarded every dataset from lab-reared fish or fish obtained through aquaria trade. Furthermore, we selected only adult individuals based on the adult body size for the species [Tsuboi et al., 2018; Froese, and Pauly, 2019]. Finally, species were included when at least ten individual measurements of body and brain size were reported. Using these rigorous selection criteria, we could include 52 species, with sample sizes ranging from 10 to 60 individuals (the exact sample size per species is reported in Table 1).

**Table 1.** Brain-body slope estimate of the 51 fish species with 95% Confidence interval.

| Species | N | $\beta$ | CI_lower | CI_upper | Species | N | $\beta$ | CI_lower | CI_upper |
| --- | --- | --- | --- | --- | --- | --- | --- | --- | --- |
| *Altolamprologus compressiceps* | 15 | 0.43 | 0.39 | 0.47 | *Neolamprologus modestus* | 10 | 0.29 | -0.02 | 0.61 |
| *Atherina presbyter* | 29 | 0.59 | 0.47 | 0.71 | *Neolamprologus pulcher* | 24 | 0.26 | 0.12 | 0.41 |
| *Bathybates fasciatus* | 11 | 0.46 | 0.4 | 0.52 | *Neolamprologus savoryi* | 10 | 0.44 | -0.21 | 1.09 |
| *Benthochromis tricoti* | 12 | 0.50 | 0.13 | 0.86 | *Neolamprologus tetracanthus* | 24 | 0.34 | 0.19 | 0.49 |
| *Chalinochromis brichardi* | 12 | 0.36 | 0.26 | 0.45 | *Ophthalmotilapia nasuta* | 17 | 0.28 | 0.12 | 0.45 |
| *Conger conger* | 42 | 0.27 | 0.23 | 0.31 | *Ophthalmotilapia ventralis* | 11 | 0.32 | -0.07 | 0.71 |
| *Coris julis* | 14 | 0.52 | 0.4 | 0.65 | *Parablennius gattorugine* | 18 | 0.51 | 0.4 | 0.62 |
| *Ctenochromis horei* | 14 | 0.46 | 0.37 | 0.56 | *Perissodus microlepis* | 10 | 0.37 | -0.01 | 0.75 |
| *Ctenolabrus rupestris* | 39 | 0.49 | 0.43 | 0.56 | *Petrochromis famula* | 12 | 0.42 | 0.25 | 0.59 |
| *Cyathopharynx furcifer* | 11 | 0.30 | 0.2 | 0.41 | *Petrochromis orthognathus* | 14 | 0.49 | 0.32 | 0.66 |
| *Cyprichromis leptosoma* | 12 | 0.55 | 0.36 | 0.74 | *Pseudosimochromis curvifrons* | 12 | 0.57 | 0.43 | 0.7 |
| *Dicentrarchus labrax* | 14 | 0.44 | 0.41 | 0.47 | *Scomber scombrus* | 11 | 0.64 | 0.5 | 0.77 |
| *Ectodus descampsii* | 14 | 0.47 | 0.17 | 0.76 | *Scorpaena porcus* | 19 | 0.38 | 0.33 | 0.43 |
| *Eretmodus cyanostictus* | 12 | 0.36 | 0.19 | 0.53 | *Serranus cabrilla* | 11 | 0.50 | 0.42 | 0.58 |
| *Gnathochromis pfefferi* | 11 | 0.71 | 0.54 | 0.88 | *Simochromis diagramma* | 13 | 0.45 | 0.35 | 0.55 |
| *Haplotaxodon microlepis* | 12 | 0.46 | 0.36 | 0.57 | *Spicara maena* | 12 | 0.49 | 0.39 | 0.59 |
| *Hippichthys cyanospilos* | 19 | 0.69 | 0.52 | 0.85 | *Spinachia spinachia* | 12 | 0.64 | 0.42 | 0.85 |
| *Hippocampus spinosissimus* | 16 | 0.47 | 0.26 | 0.69 | *Symphodus melops* | 31 | 0.49 | 0.4 | 0.58 |
| *Hippocampus trimaculatus* | 28 | 0.55 | 0.48 | 0.62 | *Syngnathoides biaculeatus* | 10 | 0.79 | 0.13 | 1.45 |
| *Julidochromis ornatus* | 23 | 0.35 | 0.21 | 0.49 | *Syngnathus schlegeli* | 60 | 0.29 | 0.19 | 0.4 |
| *Labrus bergylta* | 18 | 0.41 | 0.34 | 0.48 | *Telmatochromis temporalis* | 24 | 0.48 | 0.41 | 0.55 |
| *Lepidiolamprologus elongatus* | 15 | 0.41 | 0.35 | 0.47 | *Trematocara unimaculatum* | 10 | 0.47 | 0.23 | 0.71 |
| *Lepidiolamprologus profundicola* | 14 | 0.48 | 0.41 | 0.55 | *Tropheus moorii* | 15 | 0.39 | 0.12 | 0.66 |
| *Limnochromis staneri* | 10 | 0.41 | 0 | 0.82 | *Tylochromis polylepis* | 14 | 0.57 | 0.36 | 0.79 |
| *Lipophrys pholis* | 19 | 0.48 | 0.38 | 0.58 | *Xenotilapia melanogenys* | 15 | 0.46 | 0.23 | 0.7 |
| *Lobochilotes labiatus* | 14 | 0.44 | 0.39 | 0.48 |  |  |  |  |  |

To estimate the log brain-log body regression slope of every species from the fishes dataset, we fitted a set of Linear models (LMs with Ordinary Least Squares approach OLS). We validate these LMs by checking model assumptions like the normal distribution and homogeneity of variance of the residuals. Furthermore, we estimated the range of body sizes in a given species’ dataset for each of the 51 species by simply calculating the body size (as body mass in g) ratio of the largest (i.e., heaviest) to the smallest (i.e., lightest) individual. We then tested whether there is an effect of body size ratio on these species brain-body slopes. To do so, we fitted a phylogenetic generalised least-squares regression (PGLS) of log brain-log body slope (i.e., response variable) on log-transformed body size ratio (i.e., continuous predictor). We also explored whether sample size (log-transformed number of individuals per species) affected the estimated brain-body slopes by fitting a PGLS. The PGLS approach considers the phylogenetic relationships to produce an estimate of expected covariance in cross-species data. For this, we estimated Pagel’s λ (lambda) as the phylogenetic signal in PGLS models. Pagel’s λ is an estimate of the extent to which correlations in given traits reflect their shared evolutionary history cross-species. To calculate Pagel’s λ, we ran a maximum likelihood function for each PGLS model separately (see code for further details). For the PGLS models to meet the assumptions of normal distribution and homogeneity of variance, we had to exclude one species (*Neolamprologus multifasciatus*) from the analyses because it had a studentized residuals larger than three [Jones, and Purvis, 1997], bringing thus our sample size to 51 fish species.

We ran a phylogenetic signal analysis to test whether the Pagel’s λ of the body-brain slope is significantly different from zero (the null hypothesis). A significant λ > 0 (p ≤ 0.05) would mean that phylogeny affects the value of the estimated body-brain slopes. Furthermore, we ran a multiple comparisons analysis on the 51 fish slopes by fitting a global model for brain-body size with species as a factor. *Post hoc* tests with the Tukey correction method helped in comparing brain-body slopes between all these fish species. A detailed step-by-step code is provided as a script file that runs in the open-source statistical software R [R Core Team, 2020] (see Data and code accessibility statement).

### Case study: Brain size, body size and cognitive performance data in Labroides dimidiatus

We used published data on *Labroides dimidiatus* as a case study to test the link between brain size and body size, on the one hand, and cognitive performance and body size, on the other hand. There were no data on brain size and cognitive performance available at the individual level. All the data we use here is from fish collected on the reef around Lizard Island, Great Barrier Reef, Australia, between 2016 and 2019. Cleaner fish have a lifespan of about five years [Eckert, 1987]. They undergo a pelagic larvae phase before settling on a coral reef [Victor, 1986]. Therefore, all fish sampled in these datasets belong to the same population [see Triki et al., 2018].

#### 1. Brain size data

We compiled datasets from published studies by Triki et al. [2019a; 2020b] for brain and body size data. These two studies were conducted on wild-caught female cleaner fish *L. dimidiatus* at Lizard Island. Both studies collected fish body size as body weight to the nearest 0.01 g but differed with respect to the method used to quantify brain size: Triki et al. [2019a] used the magnetic resonance image (MRI) technic to estimate brain volume (in mm^3^) of N = 15 female cleaner fish, while Triki et al. [2020a] manually dissected and weighed the brains of N = 18 female cleaner fish to the nearest 0.1 mg (for further details, please refer to the original studies by Triki et al. [2019a; Triki et al., 2020a]). These data sets also had the volume/mass of the five major regions of fish brains, i.e., telencephalon, diencephalon, midbrain, cerebellum, and brain stem.

#### 2. Cognitive performance data

We compiled data from the study by Aellen et al. [2021 Preprint] for the cognitive performance and body size data. The dataset comprises cognitive performance of N = 69 wild-caught female cleaner fish tested in four different laboratory tasks at Lizard Island, Great Barrier Reef, in Australia. Here, we used these data to explore the relationship between body size and cognitive performance. We provide below brief descriptions for each cognitive task, but for more details, please refer to the original study by Aellen et al. [2021 Preprint]. Note that given the nature of cleaner fish feeding behaviour (i.e., removing ectoparasites off the skin of client fish), researchers often feed them in captivity with a paste of mashed prawn or a mixture of fish flakes and mashed prawn smeared as a thin layer on Plexiglas plates. Such plates with smeared food paste can be viewed as surrogates for client fish skin with ectoparasites in laboratory settings.

##### 2.1. Reversal learning task

Aellen et al. [2021 Preprint] tested fish learning and flexibility abilities in a reversal learning task. The experiment was based on a simultaneous two-choice task with Plexiglas plates with colour and shape cues, i.e., yellow triangular vs green round plates. Fish first had to learn the association between the yellow triangle and a food reward. As soon as fish had learned this association, the reward contingency was reversed, and the previously non-rewarding green plate became the one with the food reward. At this stage, fish were offered a maximum of 100 trials (i.e., ten sessions of ten trials each over five consecutive days) to learn the reversal association. The learning criterion set for this task is that each individual has to significantly (p ≤ 0.05) learn the rewarding colour above chance level (50%). That is, successful fish were those that scored either seven correct choices out of ten in three consecutive test sessions (i.e., 21/30), eight correct choices out of ten in two consecutive sessions (i.e., 16/20), or at least nine correct choices out of ten in a single session (i.e., 9/10).

##### 2.2. Detour task

The detour task aimed at estimating fish inhibitory control abilities. Here, a novel Plexiglas plate with a food reward was placed behind a transparent barrier. To access the food reward, fish had to inhibit their impulses to go straight to the visible food and instead swim around the barrier to reach the food. Bumping into the barrier, however, shows that the subject is impatient to get the food. Therefore, this measure (i.e., number of head bumps) provides an estimate of inhibitory control abilities, where fewer head bumps indicate better abilities. In total, Aellen et al. tested fish inhibitory control abilities in ten detour trials. The estimated performance was in the form of the average number of fish head bumps on the barrier until reaching the food.

##### 2.3. Numerical discrimination task

To test for fish numerical abilities, Aellen et al. [2021 Preprint] used Plexiglas plates with various numerical quantities as black squares against a white background. The number of black squares ranged from one to nine. The task consisted of presenting two Plexiglas plates at a time with two different quantities of black squares. In total, there were 20 different combinations of the nine quantities. The general rule was that, in every combination, the plate depicting the largest quantity of black squares was always the rewarding plate with food items. The performance consisted of the proportion of correct choices from 160 test trials over eight days.

##### 2.4. Feeding against preferences task

During the cleaner-client cleaning interaction, cleaner fish often prefer to bite client mucus instead of cooperating and eating ectoparasites [Grutter, and Bshary, 2003]. Such events can be reproduced in laboratory conditions with Plexiglas plates as surrogates for clients, and fish flakes and mashed prawn as surrogates for client ectoparasite and mucus, respectively [Bshary, and Grutter, 2006]. Aellen et al. [2021 Preprint] used a similar laboratory set-up to test whether cleaner fish can inhibit their food preferences for prawn and feed on fish flakes instead. A Plexiglas plate with two fish flakes items and two prawn items was presented to the fish during test trials. The plate remained in the aquarium as long as fish fed on flakes items (i.e., a less preferred food as an indicator of inhibition of preferences). Upon eating a prawn item (i.e., a highly preferred food as an indicator of non-inhibition of preferences), however, the plate withdrew from the aquarium. Thus, the optimal feeding strategy to maximise food intake is to feed first on the less preferred food (flakes) before eating one highly preferred food item (prawn). We then estimated fish performance in this task as a ratio reflecting the degree of feeding against preference. To do so, we divided the total number of fish flakes items by the total of prawn items consumed throughout the 12 test trials. Next, we fitted a linear regression model of feeding ratio on log-transformed body weight for the cognition-body slope from this data.

### Data analyses for the cleaner fish brain data

We used the cleaner fish brain data to calculate the slope of brain size (either as volume or mass) as a function of body size and the regression slope of every brain part on body size. To do so, we fitted Linear regression models LMs (with OLS) in the open-source statistical software R [R Core Team, 2020] of log-transformed brain measurement on log-transformed body mass. We had two separate sets of linear models, given that brain morphology was estimated once as a volume and once as a mass via MRI and manual dissection methods, respectively. We validated all these LMs by checking model assumptions like the normal distribution and homogeneity of variance of the residuals.

For data analyses linked to the cognitive tasks, we ran a Generalised Linear Model (GLM) with quasipoisson distribution error to estimate the regression slope of cognitive performance on log-transformed body weight (g). We ranked individual performance with respect to the performance of all tested fish. In all tasks, we ranked fish in a way that higher rank values corresponded to better performance. We used rank data as cognitive performance (i.e., response variable) in the fitted GLMs, where log-body size was the continuous predictor. We fitted GLMs with quasipoisson to solve an overdispersion issue with the Poisson distribution error.

## Results

### The intraspecific brain-body slope in fishes

We calculated intraspecific slopes in wild-caught individuals from 51 fish species (Fig. 1a). These slopes ranged from from 0.26 to 0.79 (Table 1), with a mean value of 0.46, and a median also of 0.46. Sample size (the number of individual data points per species) did negatively affect the brain-body slope estimates, though the effect size was relatively small (PGLS: N =51, β = -0.105, t-value= -2.846, *p* = 0.006, 95% CI -0.18, -0.03], R^2^= 0.13, λ = 0.55, Fig. 1b). Most importantly, we found no effect of body size ratio on the brain-body slope estimates (PGLS: N =51, β = -0.013, t-value= -0.682, *p* = 0.498, 95% CI -0.05, 0.03], λ = 0.47, Fig. 1c). Further analyses showed that there was no significant phylogenetic signal in the brain-body slopes (p-value of the likelihood ratio test = 0.16). Nevertheless, the multiple comparisons analysis of fish slopes indicated that some fish species had significantly different brain-body slopes from each other (Fig. 2).

**Figure 1.**
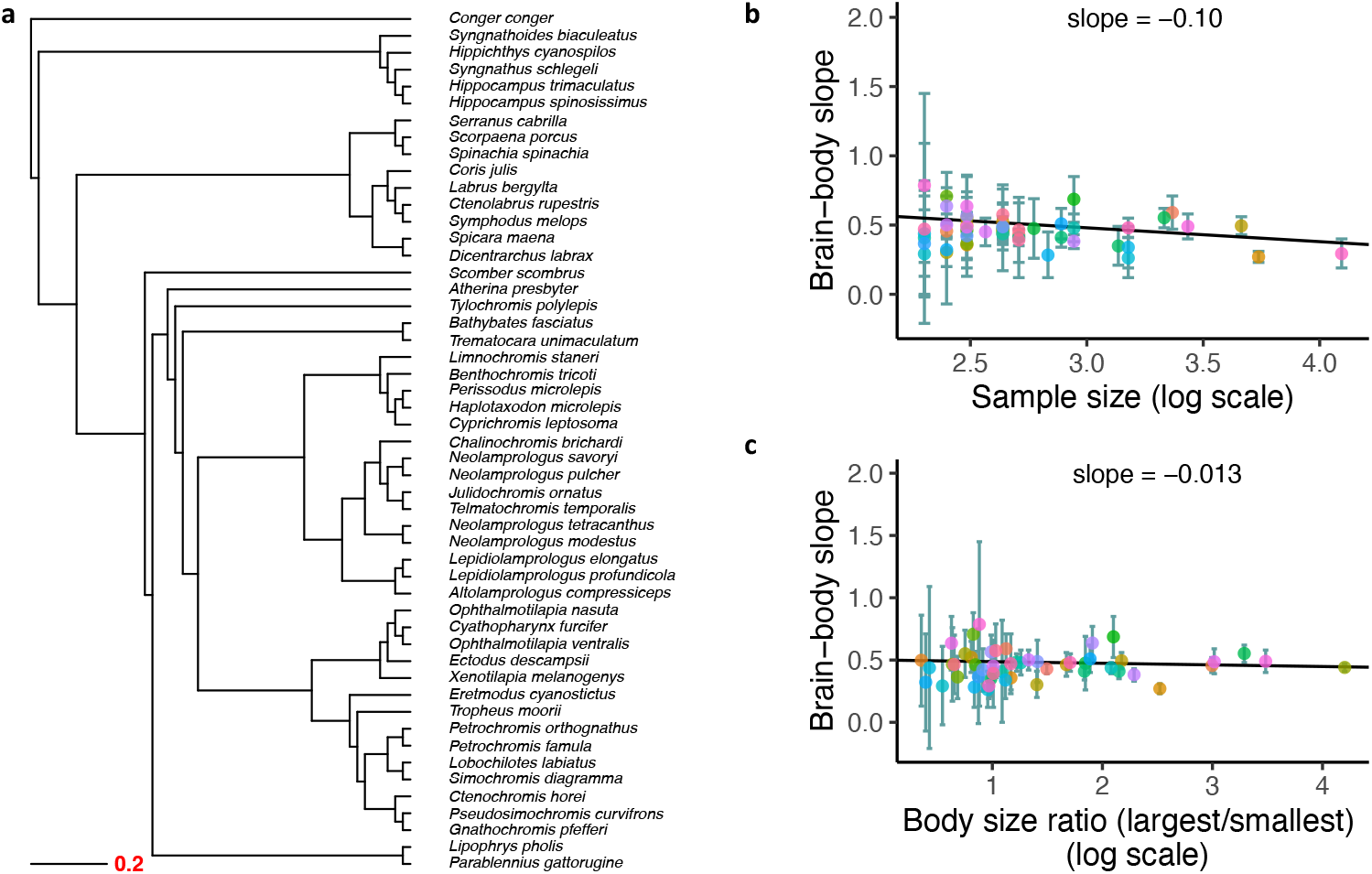
(**a**) Phylogenetic tree of the 51 fish species with a scale indicating tree distance. Relationship of brain-body slope and either (**b**) sample size used to estimate the brain-body slope, or (**c**) body size ratio. The plots show the brain-body slope estimate with 95% CI as error bars for each species. Brain and body data were compiled from the fish database FishBase [Froese, and Pauly, 2019] and from Tsuboi et al. [2018].

**Figure 2.**
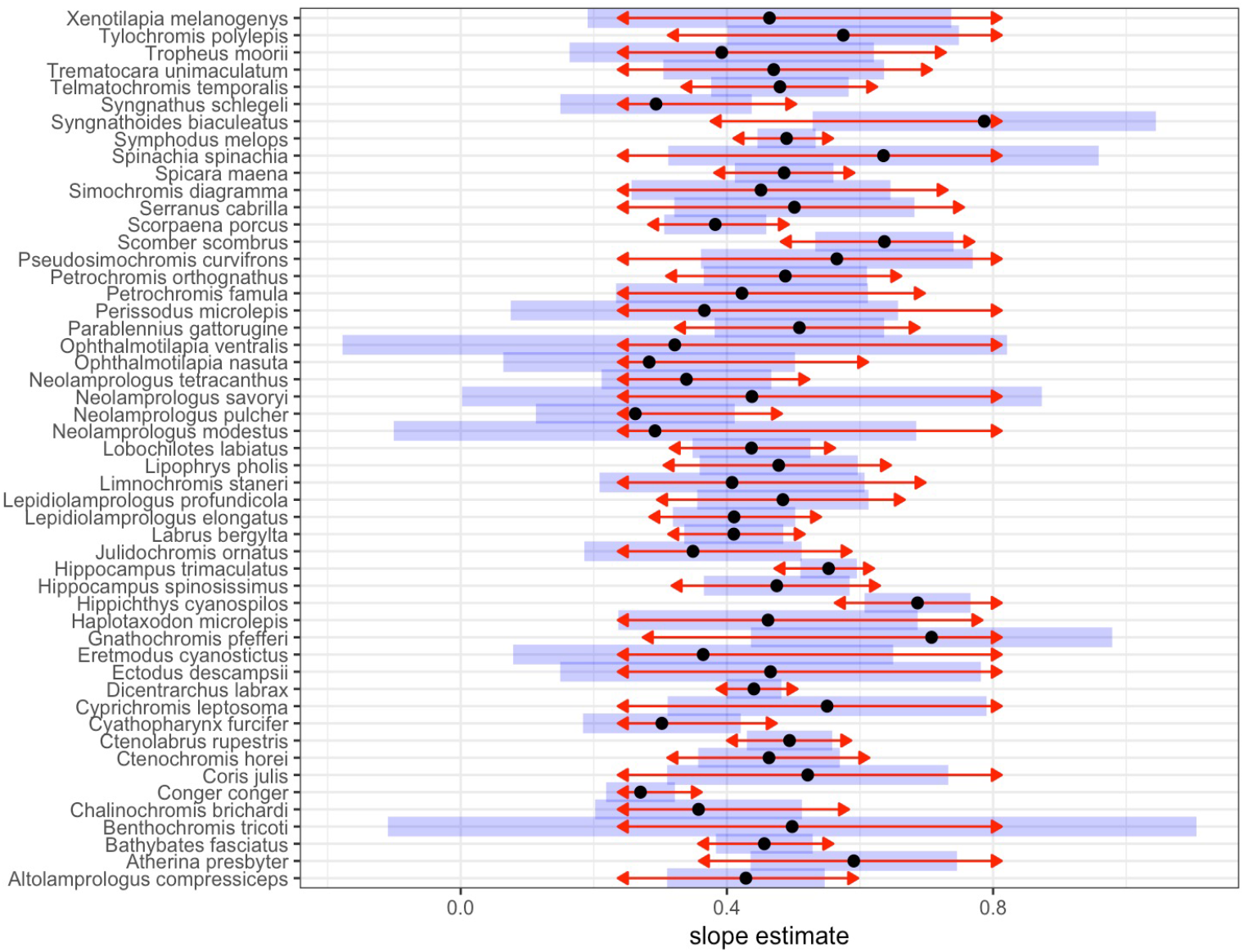
Comparison of the 51 fish slopes. Black dots are the slope estimate of log-transformed brain size on log-transformed body size. Blue bars are the 95% CI estimated from the multiple comparisons. The red arrows are indicators for the statical comparisons. If an arrow from one slope estimate overlaps with an arrow from another species, the difference is not statistically significant. We used Tukey methods to adjust for multiple comparisons with an alpha set at 0.05.

### Cleaner fish slopes

Our two datasets used to calculate the slope for the relationship between log brain size and log body size yielded values of 0.58 (MRI scans, LM: N = 15, β = 0.585, t-value= 6.133, *p* < 0.001, 95% CI [0.38, 0.79], Fig. 3a) and 0.47 (dissection data, LM: N=18, β = 0.473, t-value= 3.593, *p* = 0.002, 95% CI [0.19, 0.75], Fig. 4a), respectively. Furthermore, the regression slopes of every major brain part (i.e., telencephalon, diencephalon, midbrain, cerebellum and brain stem) on body size are reported in Table 2 and Fig. 3 & 4. The goodness of fit seems to be sensitive to the method used in estimating the size of these regions, wherein the MRI methods yielded the best estimates with a narrower 95% CI compared to the values from the manual dissection data (Figs. 3&4). Overall, brain part slopes showed some variation but were not fundamentally different from the total brain slopes. For example, the telencephalon to body size slopes of 0.54 with MRI and 0.49 with manual dissection are similar to the total brain’s slopes (Fig. 3b & 4b).

**Table 2.** Brain-body slope estimate and brain part-body slope in the cleaner fish *L. dimidiatus* with 95% Confidence interval.

| Methods | Brain measure | N | $\beta$ | CI_lower | CI_upper |
| --- | --- | --- | --- | --- | --- |
| *MRI (volume in mm<sup>3</sup>)* | Total brain size | 15 | 0.58 | 0.38 | 0.79 |
|  | Telencephalon | 15 | 0.54 | 0.23 | 0.85 |
|  | Diencephalon | 15 | 0.65 | 0.41 | 0.88 |
|  | Midbrain | 15 | 0.5 | 0.33 | 0.67 |
|  | Cerebellum | 15 | 0.64 | 0.38 | 0.89 |
|  | Brain stem | 15 | 0.62 | 0.39 | 0.85 |
| *Manual dissection (mass in mg)* | Total brain size | 18 | 0.47 | 0.19 | 0.75 |
|  | Telencephalon | 18 | 0.49 | -0.16 | 1.14 |
|  | Diencephalon | 18 | 0.62 | 0.19 | 1.05 |
|  | Midbrain | 18 | 0.48 | 0.23 | 0.74 |
|  | Cerebellum | 18 | 0.28 | -0.24 | 0.8 |
|  | Brain stem | 18 | 0.31 | -0.43 | 1.04 |

**Figure 3.**
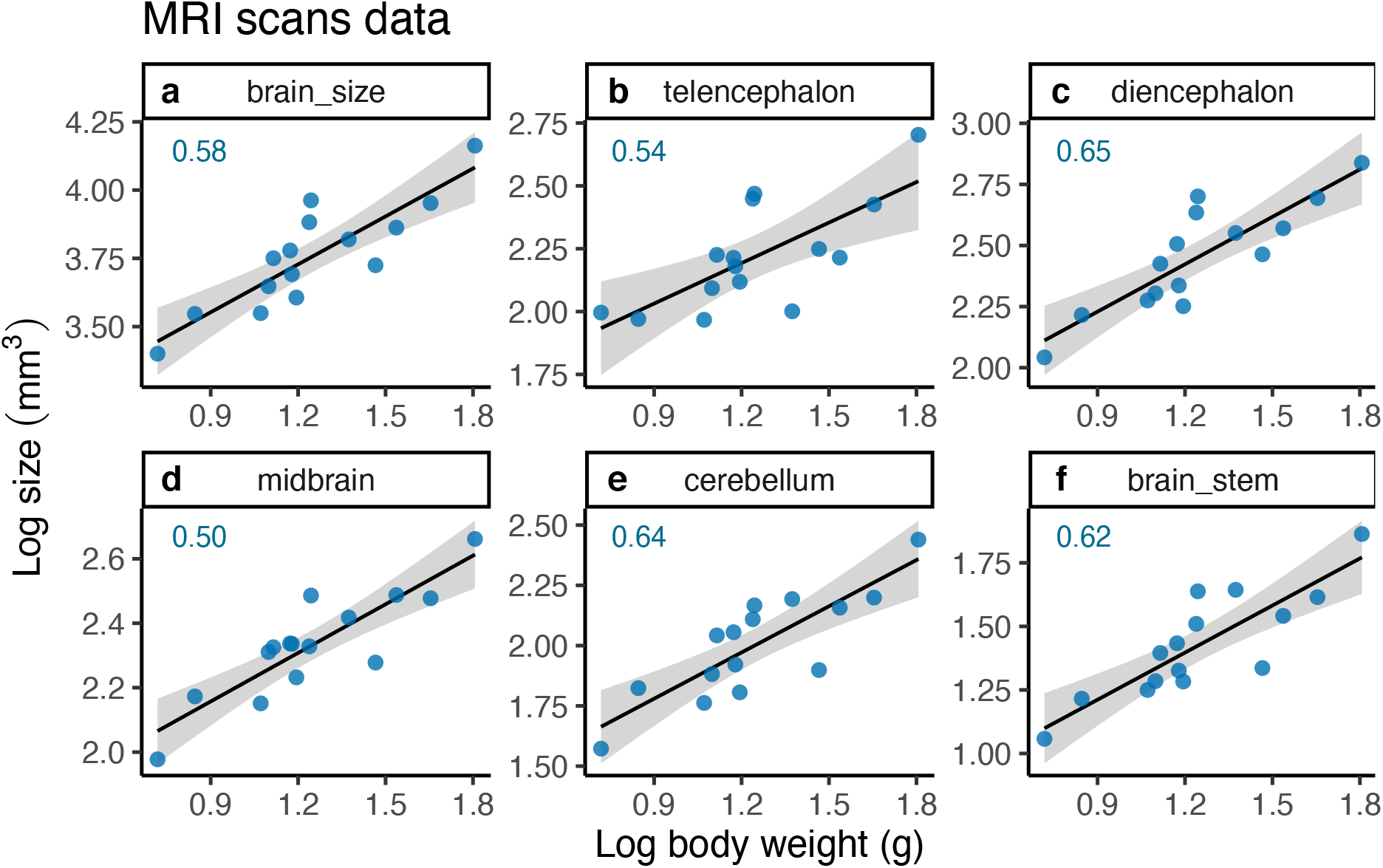
Regression relationship between (**a**) log-transformed total brain size, and (**b-f**) log-transformed major brain part sizes, and log-transformed body weight (g) in cleaner fish. Total brain and brain parts volume (mm^3^) were estimated from magnetic resonance imagery (MRI) scans (data from Triki et al. [2019a]). The shaded area around the regression lines refers to the 95% CI. Slope estimates are also depicted in the different figure panels.

**Figure 4.**
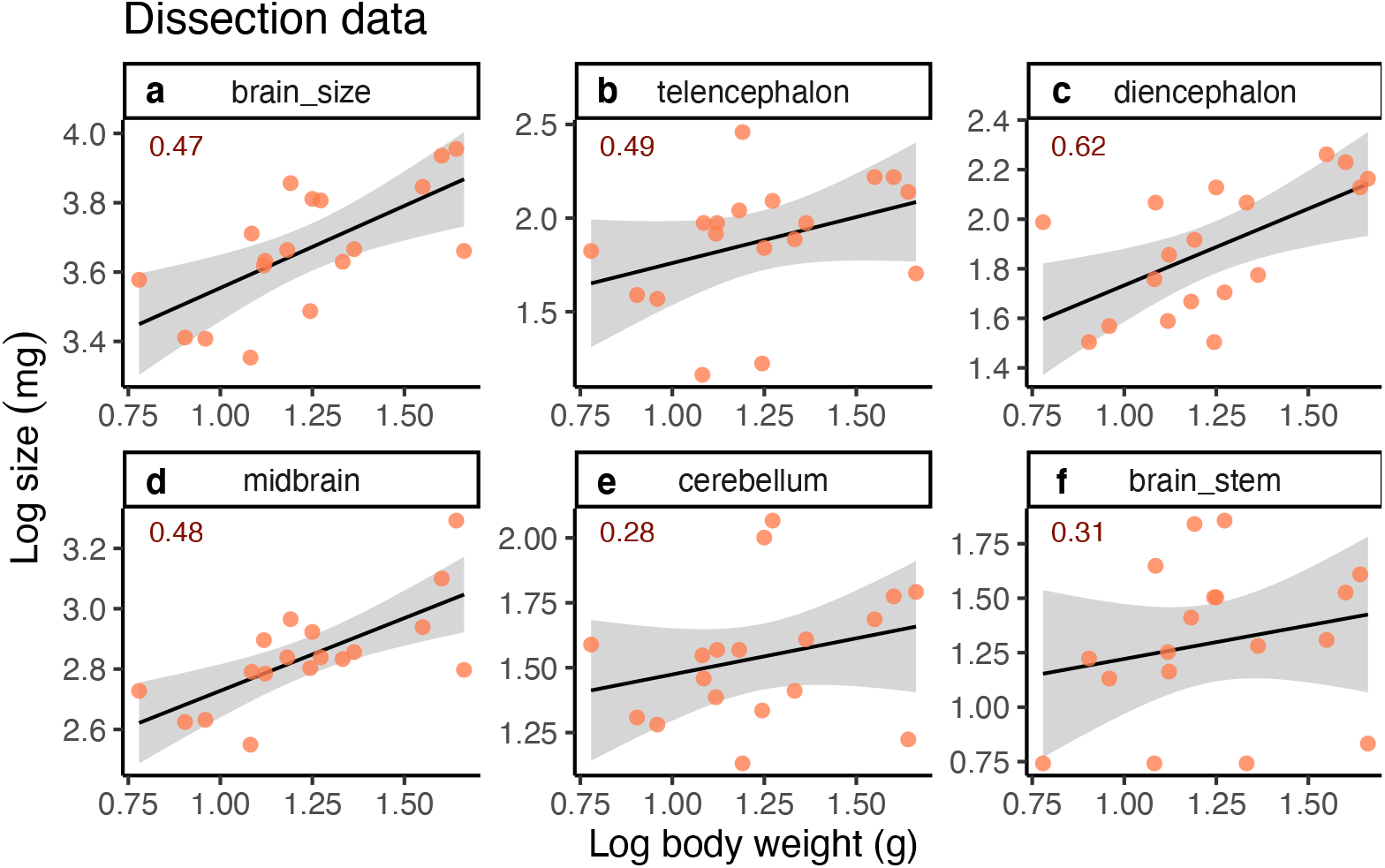
Regression relationship between (**a**) log-transformed total brain size, and (**b-f**) log-transformed major brain part size, and log-transformed body weight (g) in cleaner fish. Total brain and brain parts mass (mg) were estimated from manual dissection and weighing. The shaded area around the regression lines refers to the 95% CI. Slope estimates are also depicted in the different figure panels.

For the cognition-body slopes, there was no significant relationship between body size and performance in any of the four cognitive tasks, nor did the four slopes yield the same (positive or negative) sign: reversal learning (GLM: N=69, β = -0.400, t-value= -1.456, *p*=0.15, 95% CI [-0.94, 0.14], Fig. 5a), detour task (GLM: N=69, β = 0.180, t-value= 0.642, *p*=0.523, 95% CI [-0.37, 0.73], Fig. 5b), numerical discrimination task (GLM: N=69, β = 0.208, t-value= 0.745, *p*= 0.459, 95% CI [-0.34, 0.76], Fig. 5c), and feeding against the preferences task (GLM: N=69, β = 0.327, t-value= 1.181, *p*= 0.242, 95% CI [-0.22, 0.87], Fig. 5d).

**Figure 5.**
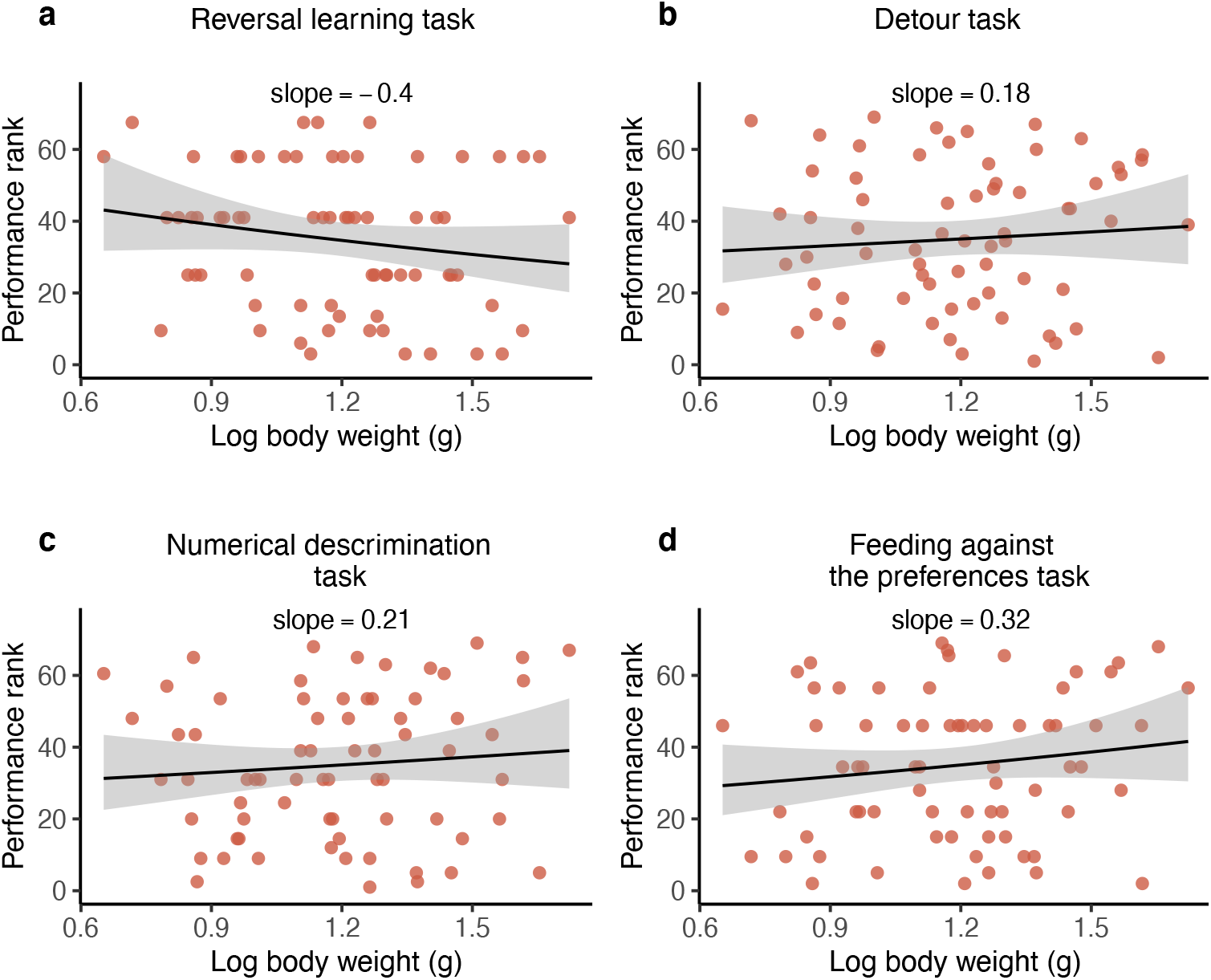
Relationship between cognitive performance and log-transformed body size (g) in cleaner fish (Data from Aellen et al. [2021 Preprint]). Regression slope and 95% CI (grey shading) of log-transformed body weight and (**a**) learning abilities in a reversal learning task; (**b**) inhibitory control abilities in a detour task; (**c**) numerical abilities in a numerical quantity discrimination task; and (**d**) inhibitory control abilities in a feeding against the preferences task. All y-axes represent performance as rank, where high values on the y-axes refer to higher performance. In all four panels, *P-values* > 0.05 were estimated from GLMs.

## Discussion

We had first asked to what extent intraspecific slopes of the relationship between body and brain measures are higher in fishes than in mammals and to what extent variation in adult body size contributes to the differences. Because the results obtained by Tsuboi et al. [2018] might have been biased, we repeated their analysis with a restrictive data set. This exercise closely replicated their findings and revealed no apparent effect of adult body size variation in fishes. We then asked whether intraspecific slopes provide a useful hypothesis regarding cognitive equivalence within a fish species, potentially between fish species as well. The key result in a case study on cleaner fish *L. dimidiatus* was that individual performance in four cognitive tasks was independent of body size. Consequently, the observed brain-body slopes (Fig. 3&4) of *L. dimidiatus* – which are rather average for a fish species – apparently reflect correctly how much brain size must be increased when the body grows to deal with somatic maintenance processes without affecting cognitive performance in this species. Below we discuss how these findings can have major implications for our understanding of the evolution of cognitive performance within vertebrates.

### Brain-body slopes in fishes

Restricting data to species that included only wild-caught specimens and a minimal sample size of ten individuals, we found an average intraspecific brain-body slope of 0.46 for Actinopterygii, compared to 0.44 in Tsuboi et al. [2018]. The small negative effect of sample size on the reported slope is difficult to interpret as there is no obvious publication bias in our data: body and brain weight data are not published as a function of brain-body slopes. Importantly for us, the effect is small and hence does not affect the general conclusions presented above. Furthermore, the absence of an effect of adult size variation on the slope estimate indicates that there is at most a minor effect of measurement error on slope estimates. Thus, our reevaluation of intraspecific brain-body slopes in fishes revealed the robustness of the slope estimates presented by Tsuboi et al. [2018] based on their much larger but less conservative data set. This finding, therefore, corroborates that fish brain-body slopes (and probably those of other ectotherms, too) are far steeper than those of birds and mammals [see also van Schaik et al. 2021]. There was an important variation of slope estimates among fish species, with minor effects of phylogenetic proximity. This suggests that targeted studies on specific fish species need to measure the slope for the species in question to generate predictions regarding the cognitive performance of individuals of this species. In our view, the average slope estimate of 0.46 is useful only for big comparative questions regarding vertebrate brains. Therefore, a major question arising is whether intraspecific slopes in fishes represent the slope of cognitive equivalence, as they do in mammals.

Our study also confirms the absence of a taxon-level effect in Actinopterygii reported by Tsuboi et al. [2018], with the brain-body slopes for class, order, family, genus and species being 0.50 ± 0.01, 0.51 ± 0.03, 0.49 ± 0.02, 0.50 ± 0.03, and 0.44 ± 0.02 (mean ± SE), respectively. This confirmation for fishes contrasts with the taxon-level effect in mammals found by Tsuobi et al. [2018] and recently confirmed for primates [van Schaik et al., 2021]. This finding, too, raises a major question concerning the use of EQs in fishes. We will discuss these two major questions in turn.

### A case study on brain-body slopes and cognitive performance in cleaner fish

We used existing data sets on *L. dimidiatus* to obtain the first preliminary results on the correlations between adult body size and cognitive performance and between adult body size and brain (region) size. Even though the two brain slope data sets available for *L. dimidiatus* yielded much higher values than birds and mammals, we did not find any systematic relationships between an individual’s body size and its performance in four different cognitive tasks. The tasks we used here tested various important cognitive domains, such as flexibility in the reversal learning task, self-control in the detour task and numbering skills in the numerical discrimination task [Burkart et al., 2017; Aellen et al., 2021 Preprint]. The fourth task – the ability to feed against preference – is of such high ecological relevance for cleaner fish in their interactions with client fish [Grutter, and Bshary, 2003] that tested cleaner fish must have had abundant prior knowledge about the consequences of eating highly preferred food (i.e., such as biting off client’s mucus). More precisely, with an estimated number of cleaning interactions for a single cleaner ranging between 800 and 3,000 per day [Grutter, 1995; Wismer et al., 2014; Triki et al., 2018; Triki et al., 2019b], all cleaner fish subjects must have had experienced in hundreds of thousands of interactions that eating parasites off clients has no negative effects while eating client mucus often leads to the client terminating the interaction [Bshary, and Grutter, 2002]. Thus, performance in our tasks was unlikely to be affected by experience. As the tasks did not address particular challenges in perception and execution of decisions but learning and decision-making, total brain size in the analyses could have been misleading. However, our analyses on the slopes of major brain regions revealed strong positive correlations between all slopes, especially with a more measurement technic like MRI. Thus, our conclusions regarding the slope of cognitive equivalence do not change much whether we take the total brain or the telencephalon, for example.

As we do not have brain data and performance data from the same fish, we could not evaluate whether individuals with a brain size above/below the slope perform above/below average. However, other studies reported that brain size or telencephalon size indeed predicts positively cognitive abilities in guppies in a reversal learning and a detour task [Buechel et al., 2018] [Triki et al., 2021b]. Thus, taken together, while we concede that our data set is preliminary, the current evidence conforms with the hypothesis that the brain-body slope also represents the slope yielding cognitive equivalence in cleaner fish.

### Implications for ectotherm-endotherm differences

In combination with a recent study on brain-body slopes in primates [van Schaik et al., 2021], the results presented here have fundamental implications for predictions on the relationship between absolute brain size, brain size relative to body size and cognitive abilities across major vertebrate lineages. The primate study, which has taken extra care to reduce the effects of any measurement errors, found that intraspecific cognitive equivalence is achieved with a brain-body slope of 0.25 to 0.30 [van Schaik et al., 2021]. This means that if the results for cleaner fish are representative for Actinopterygii, bony fishes need, on average, an increase in (log) brain size with increasing (log) body size that is almost twice as steep to maintain cognitive equivalence than primates do. By extension, we hypothesise that ectotherm vertebrates systematically need steeper brain-body slopes than endotherm vertebrates do to achieve cognitive equivalence. This is because a strong taxon-level effect has been shown for mammals (0.25-0.3 for species according to our estimates, 0.59 for class) and birds (ca 0.15 for species, which is not corrected for measurement error and thus too low, 0.57 for class) [Tsuboi et al., 2018], while it is absent or considerably smaller in the major ectotherm vertebrate lineages (Chondrichthyes: 0.5 for species, 0.41 for class; amphibians: 0.36 for species, 0.46 for class; non-avian reptiles: 0.41 for species, 0.56 for the class). In conclusion, our preliminary tests suggest that among both ectotherms [this study] and endotherms [van Schaik et al., 2021] cognitive performance is unrelated to body size within species, even though the brain-body slope in mammals is around 0.25 [van Schaik et al., 2021] and in fishes is around 0.5. If further intraspecific tests validate this assumption, this points at fundamental differences with respect to how brain organisation translates into cognitive performance between mammals and fishes, which warrant a more detailed examination.

The use of EQ as a predictor for cognitive equivalence across species has been criticised for various reasons. Some authors have proposed that absolute brain size is a better predictor than relative brain size [Rensch, 1973; Deaner et al., 2006]. Others have pointed out that some brain areas affect cognitive performance much more strongly than others [Dunbar, 1992; Healy, and Rowe, 2007; Sherry et al., 1989; Lefebvre et al., 1998]. In addition, more recent analyses show that neuron cell densities may differ between major vertebrate clades like birds and mammals and between major mammalian lineages [Herculano-Houzel, 2017], which may impose important limitations on the value of gross comparisons of brain size between more distantly related species. These problems have led to the proposal that a bottom-up approach, which focuses on comparisons between closely related species, should complement the dominating literature on large-scale comparisons [Logan et al., 2018]. Finally, van Schaik et al. [2021] had criticised the EQ concept largely because of the taxon-level effects reported for endothermic vertebrates. Indeed, the more we will eventually know about the various features of brain morphology, the better we will link them to variation in cognitive abilities. Our statistical testing of slope values also revealed significant variation among fish species, and a future challenge will be to search for potential ecological correlates of that variation. Consequently, the use of the mean intraspecific slope as a predictor for cognitive equivalence in between-species comparisons may be the best predictor we currently have, even if it will introduce large amounts of unexplained variance. Nevertheless, based on the preliminary results presented in our study, it currently appears that EQ may work far better well as a gross estimate of overall cognitive performance in ectothermic vertebrates than in endothermic vertebrates, for which its use should be discouraged [van Schaik et al. 2021]. At least the cleaner fish data suggest that large-scale interspecific comparisons may be meaningful using EQ, i.e. without more detailed knowledge on brain (part) neuron densities.

## Ethics note

There is no ethics for this present study because all data were compiled from published studies by Triki et al. [2019a; 2020b; 2020a] and Aellen et al. [2021 Preprint], and from FishBase [Froese, and Pauly, 2019].

## Acknowledgements

We thank Niclas Kolm and Tsuboi Masahito for their comments on an earlier version of the manuscript.

## Data and code accessibility

Data and code for statistical analysis and figures are accessible at the public repository Figshare. DOI: 10.6084/m9.figshare.8867660

## Authors’ contributions

CvS conceived the cognitive equivalence idea. ZT compiled the data from the different sources, analysed the data and generated the figures. ZT collected the cleaner fish brain size data. MA collected the cognitive performance data. ZT, CvS and RB wrote the manuscript. All authors approved the final version of the manuscript. A preprint version of this article is available on bioRxiv [Triki et al., 2021a, preprint].

